# HLH-30/TFEB rewires the chaperone network to promote proteostasis under conditions of Coenzyme A and Iron-Sulfur Cluster Deficiency

**DOI:** 10.1101/2024.06.05.597553

**Authors:** Rewayd Shalash, Mor Levi-Ferber, Henrik von Chrzanowski, Mohammad Khaled Atrash, Yaron Shav-Tal, Sivan Henis-Korenblit

**Affiliations:** The Mina & Everard Goodman Faculty of Life Sciences, Bar-Ilan University, Ramat-Gan 52900, Israel; The Mina and Everard Goodman Faculty of Life Sciences and Institute of Nanotechnology and Advanced Materials (BINA), Bar-Ilan University, Ramat-Gan 52900, Israel

**Keywords:** Proteostasis, protein quality control, Coenzyme A, pantothenate kinase, Iron-Sulfur clusters, TFEB, HLH-30, chaperones, *C*. *elegans*

## Abstract

The maintenance of a properly folded proteome is critical for cellular function and organismal health, and its age-dependent collapse is associated with a wide range of diseases. Here, we find that despite the central role of Coenzyme A as a molecular cofactor in hundreds of cellular reactions, limiting Coenzyme A levels in *C. elegans* and in human cells, by inhibiting the conserved pantothenate kinase, promotes proteostasis. Impairment of the cytosolic iron-sulfur clusters formation pathway, which depends on Coenzyme A, similarly promotes proteostasis and acts in the same pathway. Proteostasis improvement by Coenzyme A/iron-sulfur cluster deficiencies are dependent on the conserved HLH-30/TFEB transcription factor. Strikingly, under these conditions, HLH-30 promotes proteostasis by potentiating the expression of select chaperone genes providing a chaperone-mediated proteostasis shield, rather than by its established role as an autophagy and lysosome biogenesis promoting factor. This reflects the versatile nature of this conserved transcription factor, that can transcriptionally activate a wide range of protein quality control mechanisms, including chaperones and stress response genes alongside autophagy and lysosome biogenesis genes. These results highlight TFEB as a key proteostasis-promoting transcription factor and underscore it and its upstream regulators as potential therapeutic targets in proteostasis-related diseases.

## Introduction

The maintenance of a properly folded proteome is critical for cellular function and organismal health (Labbadia & Morimoto 2015; Meller & Shalgi 2021). High abundance of misfolded proteins drives the formation of insoluble protein aggregates that accumulate within the cell or in its vicinity, where they induce stress responses, jeopardize physiological cellular function, and lead to the development of proteinopathies, many of which are age-dependent neurodegenerative diseases affecting the elderly population (Paulson 1999; Gallotta et al. 2020; Hoppe & Cohen 2020). This underscores the importance of identifying new pathways and mechanisms that promote proteostasis in the cell and organism levels.

To counteract and minimize the damage inflicted by misfolded proteins and maintain proteostasis, a highly conserved protein quality control (PQC) system, that repairs or removes damaged proteins, has evolved (Jayaraj et al. 2020). Cellular chaperones and their co-chaperones are central components of the PQC network, and they are implicated in many aspects of protein quality control, including de novo folding of nascent polypeptides, unfolding and reactivation of denatured proteins, and controlled degradation, aggregation, or extrusion of terminally misfolded proteins (Jayaraj et al. 2020; Wentink & Rosenzweig 2023; Reinle et al. 2022). A series of dedicated stress response pathways allow the PQC system to adapt and expend according to the changing needs of the cell. This flexibility of the PQC is attributed, at least in part, to a set of stress-related transcription factors that support the expression of genes encoding protein quality control components (Pessa et al. 2024).

Coenzyme A (CoA) is a fundamental conserved cellular cofactor in hundreds of metabolic reactions (Begley et al. 2001). These include the tricarboxylic acid (TCA) cycle, mitochondrial fat metabolism and oxidation, acyl carrier protein (ACP) cofactor activation, sterol and steroid synthesis, protein acetylation and more (Leonardi et al. 2005; Strauss 2010; Pietrocola et al. 2015). CoA biosynthesis requires pantothenic acid (also known as vitamin B5), cysteine, and ATP. In this process pantothenic acid is converted into CoA by a series of five enzymatic reactions (Daugherty et al. 2002). The first step in CoA biosynthesis, which is also the rate limiting step, is the phosphorylation of pantothenic acid by pantothenate kinase (PanK) (Robishaw & Neely 1985). Mutations in the human mitochondrial PanK isoform PANK2 gene limit CoA levels and cause classical pantothenate kinase-associated neurodegeneration syndrome (PKAN) - a debilitating monogenic neurodegenerative motor-disorder characterized by movement abnormalities, vision impairment, dementia, iron accumulation in the brain, ventricular dysfunction and ultimately, premature death (Zhou et al. 2001; Shalash et al. 2021; Gregory et al. 2009; Audam et al. 2021). Similarly, mutations in the COASY gene, which acts further downstream in the CoA biosynthesis pathway, result in a related disorder known as CoA synthase protein-associated neurodegeneration (CoPAN) (Di Meo et al. 2020; Dusi et al. 2014). On the other hand, limiting CoA availability can be beneficial under some circumstances, for example in the context of a tumor, where CoA deficiency limits tumor progression (Kreuzaler et al. 2023).

Among its many roles, CoA is required for the biogenesis of Iron–sulfur (Fe/S) clusters (ISCs) (Lill & Freibert 2020). ISCs are iron and sulfur ions assembled into scaffold complexes in the form of 2Fe–2S and 4Fe–4S clusters (Shi et al. 2021). Fe-S clusters can accept or donate electrons to carry out oxidation/reduction reactions and facilitate electron transport (Rouault 2015; Webert et al. 2014). In the nucleus, Fe-S clusters are cofactors of several crucial DNA repair enzymes. All in all, ISCs are cofactors of more than 200 Fe/S proteins and are implicated in a variety of metabolic reactions (Paul & Lill 2015).

ISCs are usually assembled in the mitochondria, by a dedicated ISC machinery (Paul & Lill 2015). ISCs generated in the mitochondria can assemble with a variety of protein complexes within the mitochondria, but can also be exported to the cytoplasm and nucleus. ISC export is carried out via dedicated transporters such as the ABC transporter ABCB7, which deliver them to the cytosolic iron-sulfur protein assembly (CIA) machinery (Lill & Mühlenhoff 2008). In the cytoplasm, specialized proteins such as CIAO1 and CIA2B load the ISCs onto target cytosolic Fe–S proteins (Lill et al. 2014). Deregulation of mitochondrial and cytoplasmic ISC components and substrates have been linked to numerous human diseases (Sheftel et al. 2010; Maio & Rouault 2020).

In this work, we explored how deficiencies in CoA and ISC production affect the proteostasis network. Unexpectedly, we find that limiting CoA and cytosolic ISCs levels improve the resistance of the animals to a wide variety of proteostasis challenges. These conditions are associated with activation of the HLH-30/TFEB transcription factor, which is mostly known as a central autophagy and lysosome biogenesis promoting factor. Strikingly, we discovered that under conditions that limit CoA and/or ISC levels, HLH-30 promoted proteostasis by potentiating the expression of chaperone genes. These findings mark CoA and ISC availability as upstream regulators of the HLH-30/TFEB transcription factor and highlight this conserved transcription factor as a versatile context-dependent modulator of the PQC system.

## RESULTS

### *pnk-1* deficiency limits protein aggregation in a Poly-Q disease model

PANK deficiency is associated with neurodegenerative diseases in humans, and knockout of its homologs results in unhealthy short-lived animals in a variety of model organisms including *C. elegans* (Wu et al. 2009; Samuelson et al. 2007; Srinivasan et al. 2015; Mignani et al. 2021). Nevertheless, we found that hypomorphic *C. elegans pnk-1* animals, due to *pnk-1* RNAi treatment, or due to a deletion mutation upstream to the *pnk-1* gene that reduces *pnk-1* transcript levels by approximately 50% **(Figure S1A)**, had a normal lifespan **(Figure S1B,C and Table S1).** Hence, these *pnk-1* hypomorphic mutants can be exploited for studying the physiological implications of mild *pnk-1* deficiency.

Specifically, we explored how mild *pnk-1* deficiency affected disease-associated protein aggregation using a *C. elegans* model of Huntington-like diseases. In this model, a reporter gene (YFP) is fused to multiple copies of the amino acid Glutamine (poly-Q), inducing age-dependent protein aggregation due to the poly-Q tail (Brignull et al. 2006). We examined the effect of *pnk-1* knockdown by RNAi on the aggregation of the poly-Q_35_ or poly-Q_44_ reporter in transgenic animals expressing the transgene in their muscle or intestine, respectively. Unexpectedly, following treatment with control or *pnk-1* RNAi, we observed a significant reduction in the number of detectable foci in both muscle and intestine poly-Q reporter strains on Day 5 of adulthood (**Figure 1A, Figure S2).** Thus, in spite the fact that PANK deficiency is associated with neurodegenerative diseases in humans, and its knockout results in sick short-lived animals (Wu et al. 2009; Samuelson et al. 2007; Srinivasan et al. 2015), mild *pnk-1* silencing was beneficial under these conditions as it limited protein aggregation in this disease model.

**Figure 1:**
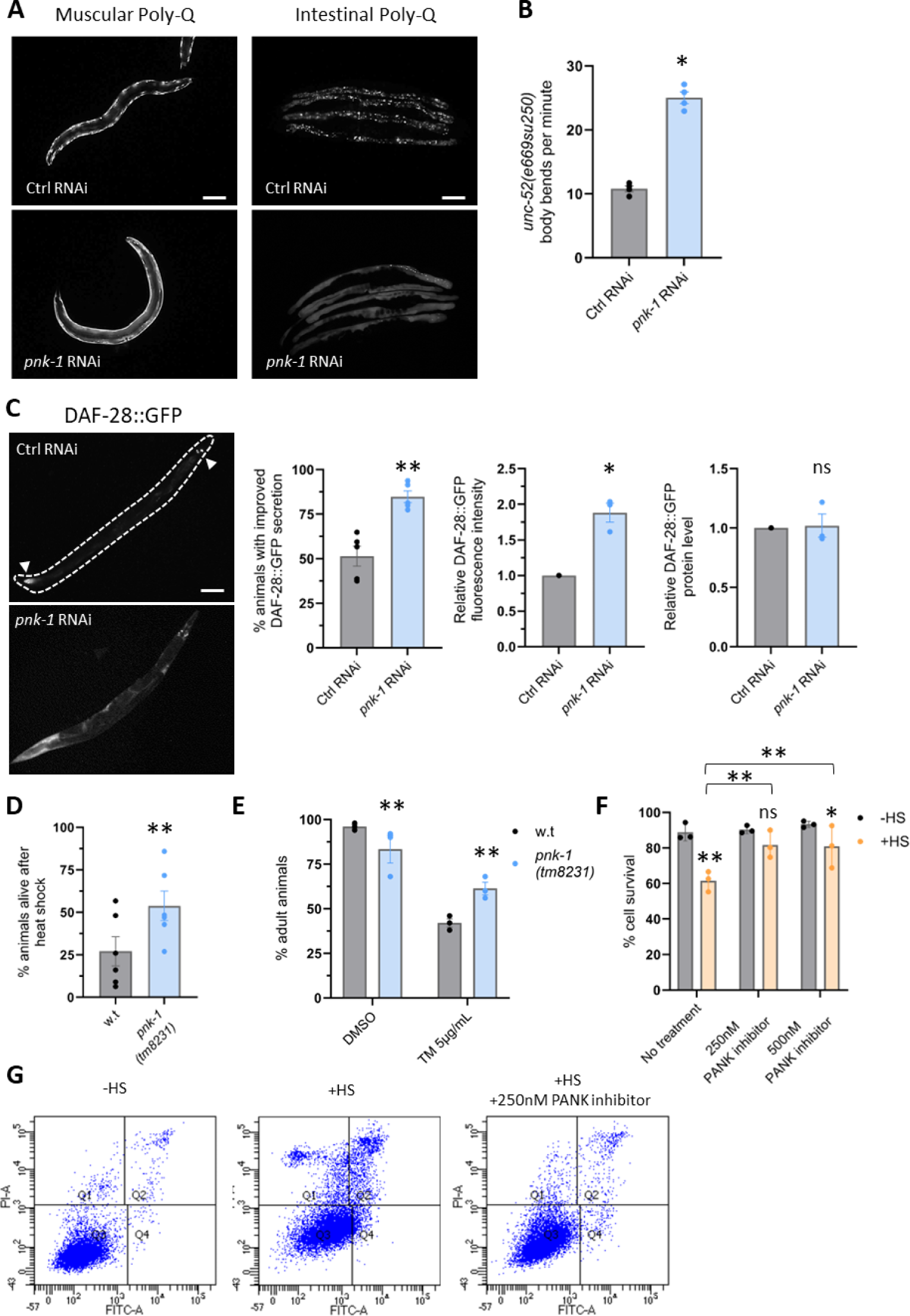
Mild CoA deficiency improves proteostasis. **(A)** *pnk-1* RNAi reduced the amount of visible aggregates in animals expressing muscular PolyQ35::YFP or intestinal PolyQ44::YFP. Animals were scored on day 5 of adulthood. See aggregate quantification in Figure S2. Scale bar: 100μm. **(B)** *pnk-1* RNAi improved thrashing of *unc-52(e669su250)* metastable mutants (N=4, n>80). **(C)** *pnk-1* RNAi improved the folding and secretion of a DAF-28::GFP reporter in *ire-1(-)* mutants. Arrow heads mark head neurons and intestinal cells that accumulate DAF-28::GFP protein in representative images of *ire-1(ok799)* mutants expressing the transgene treated with ctrl or *pnk-1* RNAi. Scale bar: 100μm. *pnk-1* RNAi increased the fraction of animals with improved DAF-28::GFP secretion (N=5, n>115), increased the overall fluorescence intensity of the DAF-28::GFP (N=3, n>75), which reflects properly folded GFP, but had no effect on total DAF-28::GFP protein level assessed by western blot (N=3), which reflects GFP protein levels irrespective to the folding state of the GFP. **(D)** *pnk-1(tm8231)* mutation improved heat-shock resistance in day 2 animals (N=6, n>200). **(E)** *pnk-1(tm8231)* mutation improved animal development in the presence of the ER stress inducer tunicamycin (N=3, n=300). **(F-G)** Annexin V/PI flow cytometry analysis of U2OS osteosarcoma cells following 2 hours of heat shock. Treatment with the PANK inhibitor increased cell resistance to heat shock (N=3). See table S1. Data are shown as mean ± standard error. N represents the number of biological repeats, n represents the number of animals analyzed per treatment or genetic background. ns, not significant; *P<0.05; **P<0.01. Statistical tests: unpaired student’s t-test **(B)**; one sample t-test values **(C)**; Cochran-Mantel-Haenszel test **(C,D,E)**, two Way Anova with Tukey’s post hoc test **(F)**. Comparisons were between w.t and *pnk-1(tm8231)* mutants per treatment **(D,E)** or relative to the corresponding control RNAi sample **(B,C)**.

### *pnk-1* deficiency promotes proteostasis in *C. elegans*

Protein aggregates are cytotoxic and highjack critical cellular components, including proteostasis-related proteins such as chaperones and co-chaperones, compromising the proteostasis network (Yue et al. 2021; Jana et al. 2000; Shirasaki et al. 2012). Hence, we hypothesized that since mild *pnk-1* deficiency limited protein aggregation, as assessed by the polyQ model, it may also lead to an overall improvement in cellular proteostasis.

Functional assays which depend on the proper folding of specific proteins offer a simple method for assessing protein folding. To investigate the impact of *pnk-1* deficiency on protein folding, we first applied a functional assay to assess the folding efficiency of a metastable protein, using a strain with a temperature-sensitive *e669su250* mutation in the *unc-52* gene, as previously described (Ben-Zvi et al. 2009). This gene encodes a critical component of the basement membrane underlying muscle cells, crucial for proper muscle assembly and muscle function. Misfolded UNC-52 impairs thrashing movement of animals in liquid. We found that treatment with *pnk-1* RNAi more than doubled the frequency of body bends per minute in *unc-52(e669su250)* mutants on day 2 of adulthood at the non-permissive temperature (**Figure 1B)**. This suggests an improvement in the folding of this metastable protein upon *pnk-1* silencing.

Another way to assess protein folding is to track proteins along the secretory pathway. To this end, we evaluated the effect of *pnk-1* deficiency on protein secretion using the DAF-28::GFP secretory model (Safra & Henis-Korenblit 2014; Safra et al. 2013). In this model, we follow the secretion of a labeled insulin protein (DAF-28::GFP) expressed in select neurons and intestinal cells in *C. elegans*. Entrapment of the protein within the producing cells indicates a secretory defect, whereas its’ successful secretion into the body cavity of the animals indicates complete processing of the protein. Furthermore, the DAF-28::GFP system allows the differential detection of the properly folded population of DAF-28::GFP protein (detected by fluorescent DAF-28::GFP) and the total population of the protein, irrespective to its folding state (detected by western blotting) (Safra et al. 2014; Safra et al. 2013). To examine whether mild *pnk-1* deficiency improves DAF-28::GFP folding and secretion, we examined DAF-28::GFP expression pattern and levels in day 3 animals with a defective ER stress response due to a mutation in the *ire-1* gene. As previously reported, these proteostasis-challenged animals fail to secrete the reporter, which remains trapped in the ER of the cells that produce it (Safra et al. 2013). We found that treatment with *pnk-1* RNAi improved the secretion of the reporter in *ire-1*-deficient animals, as reflected by the increased proportion of animals in which the DAF-28::GFP transgene was detected in their body cavity at the expense of the intestinal cells that produce it (**Figure 1C left panels)**. Furthermore, *pnk-1* RNAi significantly increased the fluorescence of the reporter but had no effect on total protein levels of the reporter (**Figure 1C right panels),** indicating an increase in the proportion of properly folded reporter. These results support the notion that mild *pnk-1* deficiency improves protein folding.

### *pnk-1* deficiency promotes stress resistance in *C. elegans* and in mammalian cells

If mild *pnk-1* deficiency improves protein folding, it should also promote animal survival upon exposure to lethal proteostasis-related stresses. To evaluate the influence of reduced *pnk-1* levels on sensitivity to proteostasis-related stresses, we conducted heat shock and tunicamycin resistance experiments with wild-type and *pnk-1*-hypomorphic animals. In the heat shock experiment, wild-type animals and *pnk-1(tm8231)* mutants were exposed to heat shock on day 2 of adulthood, and their survival was assessed 16 hours later. In the ER stress experiment, the ability of wild-type animals and *pnk-1* mutants to develop from eggs to adults in the presence of the ER stress inducer tunicamycin was examined. Consistent with its ability to improve proteostasis, the *pnk-1* mutation significantly enhanced both thermotolerance and tunicamycin resistance of the animals (**Figure 1D,E)**.

Being a key enzyme in CoA biosynthesis, PANK is conserved from bacteria to humans (Leonardi et al. 2005). Hence, we examined whether inhibition of PANK activity promotes proteostasis in human cells as well. To this end, human osteosarcoma U2OS cells were treated with PANK inhibitor and then exposed to heat shock. Cell necrosis and apoptosis were detected by Annexin V-FITC/Propidium Iodide (PI) staining. Under these stress conditions, apoptotic and necrotic cells were detected, as reflected by the increase in the proportion of the Annexin V negative/PI positive and Annexin V positive/PI positive cell subpopulations (**Figure 1G and Table S2)**. Strikingly, treating cells with a PANK inhibitor (Sharma et al. 2015) significantly limited the heat-induced cell death. Treatment with the PANK inhibitor had no affect on the viability of the cells that were not exposed to heat shock **(Figure 1 F,G and Table S2)**. These results suggest that PANK inhibition can improve proteostasis and promote heat stress resistance in human cells as well, implying that the capacity of mild *pnk-1* deficiency to promote resistance to proteostasis stresses is evolutionarily conserved.

### *pnk-1* deficiency promotes proteostasis by limiting CoA levels

PNK-1 is the rate limiting enzyme in CoA biosynthesis; hence its deficiency should limit CoA levels (Leonardi et al. 2005). To examine whether the proteostasis improvement associated with *pnk-1* deficiency is indeed due to limitting CoA levels, we performed CoA supplementation experiments. To this end, we repeated the heat shock survival assay and the DAF-28::GFP secretion assay with or without CoA supplementation. Remarkably, in both assays external supplementation with 400mM CoA negated the benefits of *pnk-1* deficiency **(Figure S3).** This indicates that proteostasis improvement because of *pnk-1* deficiency is due to limited CoA availability.

### *pnk-1* deficiency promotes proteostasis by limiting ISC availability

Many of the biochemical processes in which CoA participates in take place within the mitochondria (Leonardi et al. 2005). Thus, we examined whether impairment of mitochondrial pathways that depend on CoA promotes proteostasis similarly to *pnk-1* deficiency. To this end, we applied RNAi treatments targeting select genes comprising mitochondrial pathways which utilize CoA. These included the mitochondrial fatty acid synthesis pathway (*mecr-1, lias-1, F10G8.9*), the Krebs/TCA Cycle (*aco-2, sdhd-1, pdhb-1*), mitochondrial acyl carrier proteins (ACPs) (*T04G9.4, T28H10.1*) and the iron-sulfur cluster (ISC) biogenesis pathway (*iscu-1, nfs-1, frh-1*) (**Figure 2A).** We examined whether any of these RNAi treatments phenocopied the improvement in the secretion of the secretory reporter DAF-28::GFP, as observed with *pnk-1* RNAi treatment (**Figure 1C).** Interestingly, silencing of *mecr-1*, *lias-1* and *F10G8.9* genes implicated in mitochondrial fatty acid synthesis and silencing of *aco-2*, *sdhd-1* and *pdhb-1* genes implicated in the Krebs/TCA Cycle did not improve, and in most cases further interfered, with the secretion of the reporter (**Figure 2B,C).** In contrast, silencing of *T04G9.4* and *T28H10.1* genes encoding mitochondrial acyl carrier proteins (ACPs) and silencing of the iron-sulfur cluster (ISC) formation genes *iscu-1*, *nfs-1* and *frh-1* enhanced the secretion of the reporter (**Figure 2D,E)**. Since ACPs are also an essential subunit which stabilizes the eukaryotic Fe-S biogenesis complex (Van Vranken et al. 2016; Braymer & Lill 2017), it is likely that ACPs and ISC formation genes affect proteostasis by a common mechanism.

**Figure 2:**
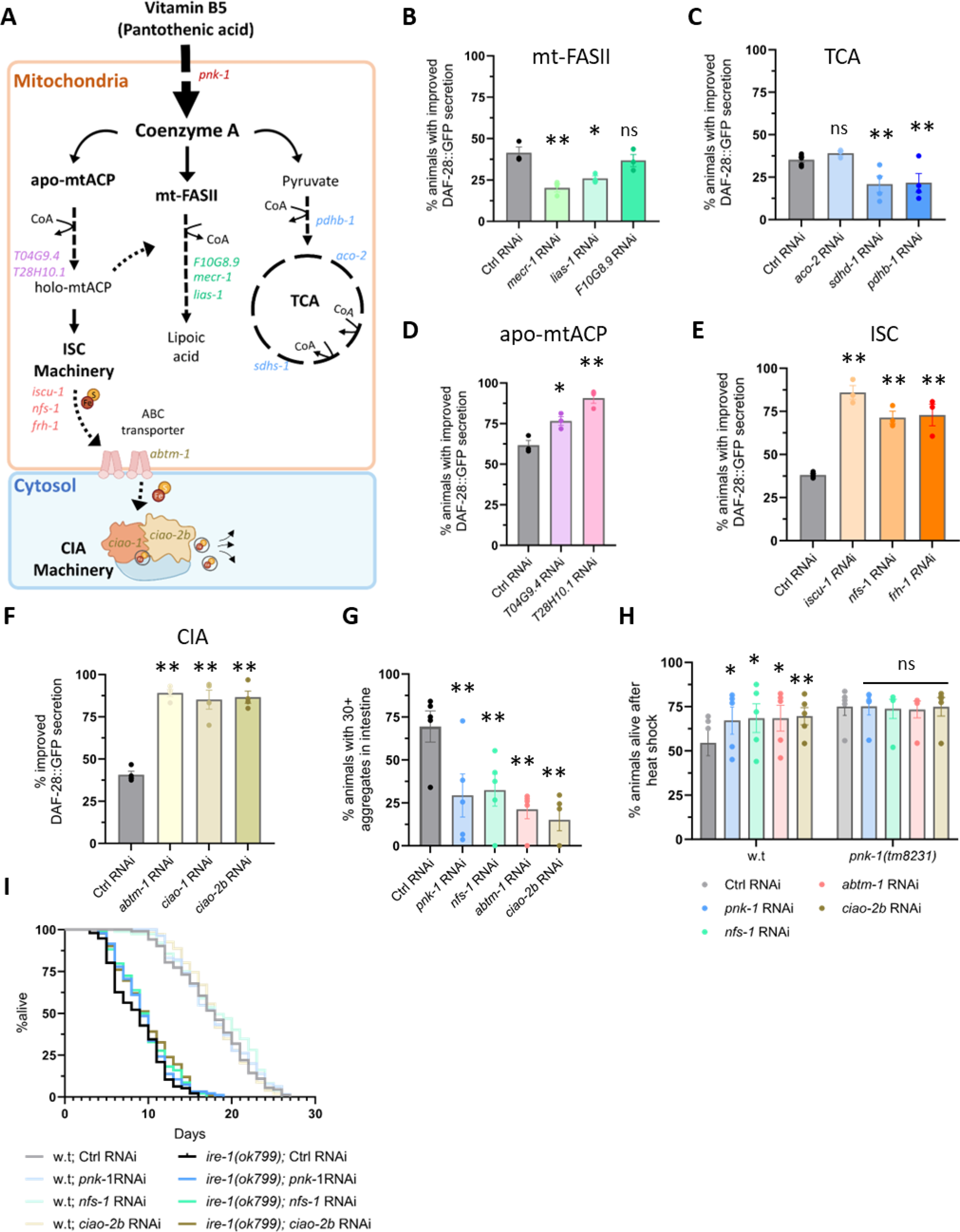
CoA and ISC deficiency act in the same pathway to promote proteostasis. **(A)** Schematic overview of the central biochemical mitochondrial pathways involving CoA including the mitochondrial fatty-acid biosynthesis pathway (mt-FASII, green), the tricarboxylic acid cycle (TCA/Krebs cycle, blue), mitochondrial acyl carrier proteins (mtACP, pink), mitochondrial iron sulfur clusters biogenesis (ISC, orange), cytosolic iron sulfur cluster assembly (CIA, yellow). **(B-F)** RNAi knockdown of genes related to mt-FASII **(B)** and TCA **(C)** pathways did not improve DAF-28::GFP secretion in day 3 *ire-1(ok799)* animals. RNAi knockdown of genes related to ACP -acyl carrier protein **(D)**, iron sulfur clusters (ISC) **(E)** and cytosolic iron sulfur cluster assembly (CIA) **(F)** improved DAF-28::GFP secretion in the *ire-1(ok799)* animals. (N≥3, n>90). See Figure S4A for representative images. **(G)** RNAi knockdown of *pnk-1* or ISC related genes reduced the number of aggregates in adult day 5 animals expressing intestinal YFP::PolyQ44 (N=5, n>150). See **Figure S4B** for representative images. **(H)** ISC related genes RNAi increased heat-shock resistance of Day 2 w.t animals but did not further increase the heat resistance of *pnk-1(tm8231)* mutants (N=5, n>140). **(I)** Representative lifespan assay of wild-type and *ire-1(-)* mutants (N=3). RNAi knockdown of *pnk-1* or ISC related genes did not extend the lifespan of wild-type or *ire-1(ok799)* animals, besides *ciao-2b* RNAi, which extended the lifespan of *ire-1(-)* mutants by 15% on average. (N=3). Data are shown as mean ± standard error. N represents the number of biological repeats, n represents the number of animals analyzed per RNAi condition. ns, not significant; *P<0.05; **P<0.01. Statistical tests: Cochran-Mantel-Haenszel test followed by FDR correction compared to the corresponding control RNAi sample **(B-H)**.

ISC are generated within the mitochondria by the mitochondrial ISC complex and are subsequently either utilized within the mitochondria or transported to the cytoplasm (Lill & Freibert 2020). In the cytoplasm, a protein complex known as the cytosolic iron-sulfur cluster machinery assembly (CIA) loads these clusters onto carrier proteins, which transfer them to target proteins outside of the mitochondria (Paul & Lill 2015). Therefore, we explored whether the improvement in proteostasis is also observed upon disruption of the CIA complex (perturbing iron-sulfur dependent activities in the cytoplasm, but not affecting those in the mitochondria). To this end, we monitored DAF-28::GFP secretion upon RNAi silencing of the Fe-S cluster transporter *abtm-1*, that facilitates cluster transfer from mitochondria to the cytoplasm, or silencing of the *ciao-1*, *ciao-2b* genes, which encode components of the cytoplasmic ISC machinery. Knockdown of all genes implicated in cytoplasmic iron-sulfur cluster metabolism improved the secretion of the DAF-28::GFP reporter (**Figure 2F,S4A).** Similarly, silencing of ISC-related genes, be it in the mitochondria or in the cytoplasm, reduced the aggregation of the poly-Q_44_ intestinal reporter (**Figure 2G,S4B)** and improved heat shock survival, similarly to *pnk-1* RNAi treatment (**Figure 2H)**. Thus, deficiency in mitochondria or cytoplasmic ISCs is sufficient for proteostasis improvement.

Finally, since CoA is required for ISC biogenesis (Shi et al. 2021) and since deficiencies in either *pnk-1* or ISC biogenesis components, in the mitochondria or in the cytosol, improved proteostasis, we hypothesized that they might operate within the same genetic pathway. To test this hypothesis, we conducted an epistasis experiment, treating *pnk-1(tm8231)* mutants with RNAi targeting ISC/CIA genes and compared their resistance to heat shock. We found that whereas these RNAi treatments increased heat shock survival of wild-type animals (**Figure 2H, left panel)**, they did not further increase heat shock survival of *pnk-1(tm8231)* mutants (**Figure 2H, right panel).** The lack of an additive effect between *pnk-1* deficiency and ISC biogenesis deficiencies supports the hypothesis that they promote proteostasis by a common mechanism. Since CoA is required for effective ISC biogenesis, these results imply that *pnk-1* deficiency promotes proteostasis by limiting ISC biogenesis, and specifically ISC availability in the cytoplasm.

### *pnk-1* and ISC deficiencies do not extend lifespan

The loss of proteostasis is a fundamental aspect of the aging process, stemming from the malfunction of the cells’ protein stress responses and the consequential age-dependent accumulation of misfolded proteins (Hipp et al. 2019). Since a reduction in CoA synthesis and ISC biogenesis significantly improved proteostasis under various stress conditions (**Figure 1,2)**, and since no lifespan benefit was observed in *pnk-1* deficient animals under normal growth conditions **(Figure S1B,C)**, we postulated that a lifespan benefit might be detected under proteostasis stress conditions. To this end, we examined how *pnk-1* deficiency affected the lifespan of animals with a perturbed ER stress response due to a mutation in the *ire-1* gene. Under these stress conditions, no consistent extension of lifespan was observed upon *pnk-1* or *nfs-1* RNAi treatment (**Figure 2I)**. A lifespan extension of 14-18% was observed upon treatment with RNAi targeting the *ciao-2b* gene, resulting in the dysfunction of the CIA complex (**Figure 2I and Table S1)**. We note that a small lifespan extension has also been reported upon RNAi targeting the mitochondrial iron-sulfur cluster assembly protein component *iscu-1* (Sheng et al. 2021). Thus, whereas mild impairment of *pnk-1* and the ISC biosynthetic pathways consistently improved proteostasis, this was not coupled to lifespan extension.

### *pnk-1* mild deficiency improves proteostasis in a chaperone dependent proteosome/lysosome independent manner

The two major protein quality control mechanisms for improving proteostasis involve protein refolding and clearance of misfolded proteins (Chen et al. 2011; Meléndez & Levine 2009). As such, we examined whether the proteostasis enhancement resulting from CoA limitation is dependent on the main protein degradation machineries of the cell (the proteasome and the lysosome) and whether it depends on cellular chaperones (which are critical for protein refolding, but also for protein disaggregation and protein degradation).

To examine the contribution of the proteasome to the proteostasis of *pnk-1* deficient animals, we tested whether the improved heat shock resistance induced by *pnk-1* deficiency and the improved DAF-28::GFP secretion in *ire-1*-deficient animals are proteosome dependent. Specifically, DAF-28::GFP protein secretion and heat shock resistance of *pnk-1* mutants was compared between animals treated with control RNAi or with RNAi targeting the proteasome regulatory subunit *rpn-6.1* starting at the L4 stage (earlier treatment with *rpn-6.1* RNAi was lethal to the animals, and demonstrated the efficacy of this RNAi treatment). Strikingly, *rpn-6*.1 RNAi treatment did not affect DAF-28::GFP secretion or heat shock resistance of wild-type or *pnk-1* mutant animals (**Figure 3A, B).** This indicates that *pnk-1* deficiency improves ER secretory function independent of the proteosome.

**Figure 3:**
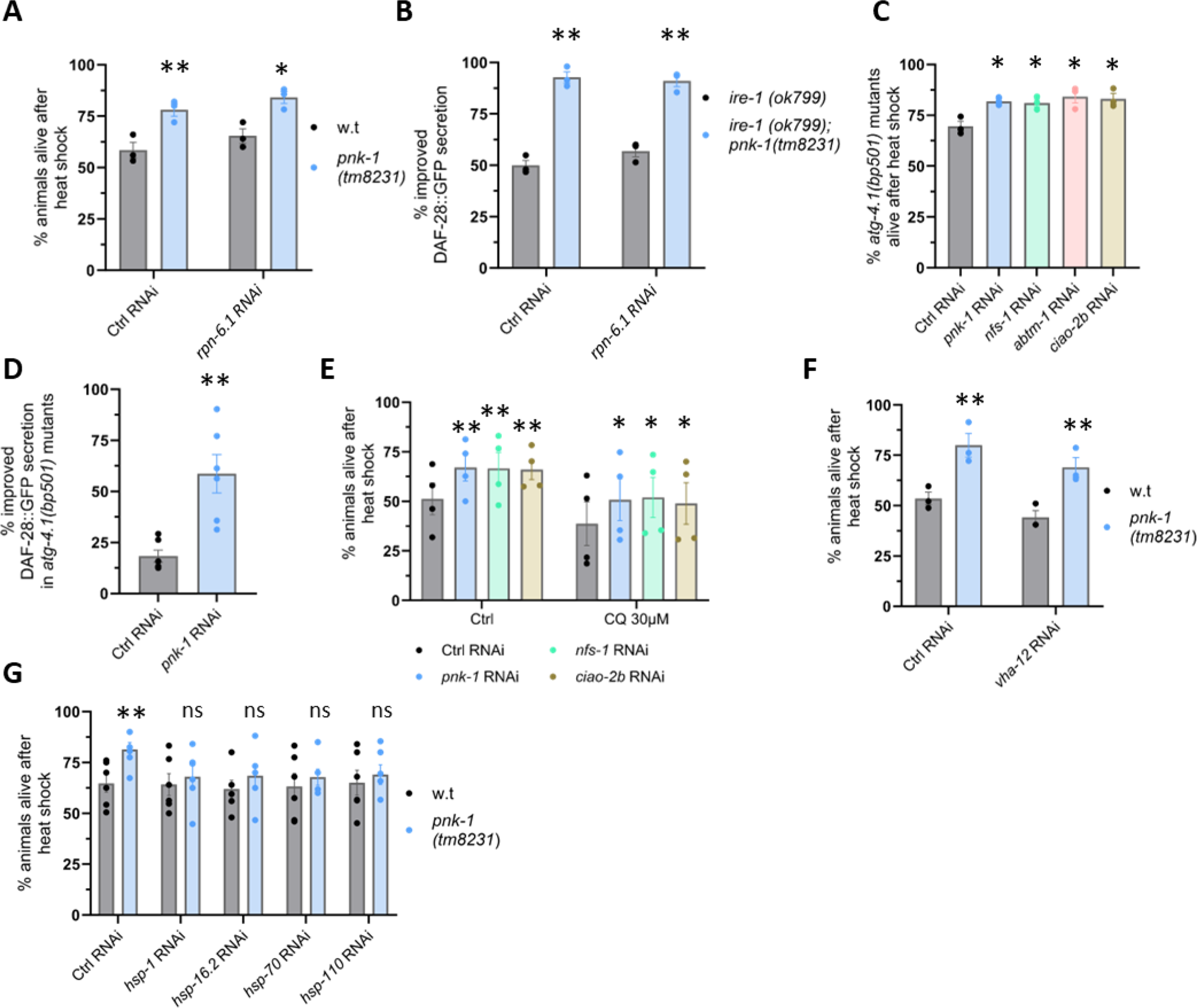
CoA and ISC limitation improved-proteostasis depends on protein chaperones. **(A-B)** Inhibition of proteasome function by *rpn-6* RNAi treatment in adult animals did not affect heat-shock resistance of *pnk-1(tm8231)* animals at day 2 of adulthood (N=3, n>80) **(A)** nor did it affect DAF-28::GFP secretion improvement by *pnk-1(tm8231)* in *ire-1(ok799)* day 3 mutants (N=3, n>70**) (B)**. **(C,E)** *pnk-1* and ISC related gene inhibition improved heat-shock resistance independent of autophagy (N=3, n>90) **(C)** and independent of chloroquine (CQ) treatment (N=4, n>160) **(E)**. **(D)** *pnk-1* RNAi improved DAF-28::GFP secretion in day 3 *atg-4.1(bp501)* mutants (N=6, n>144). **(F)** Inhibition lysosome function by *vha-12* RNAi treatment in adult animals did not affect heat-shock resistance of *pnk-1(tm8231)* animals (N=3, n>105). **(G)** RNAi knockdown of select cytosolic chaperone genes abolishes the beneficial effects of *pnk-1* deficiency on heat resistance (N=6, n>150). Data are shown as mean ± standard error. N represents the number of biological repeats, n represents the number of animals analyzed per treatment or genetic background. ns, not significant; *P<0.05; **P<0.01. Statistical test: Cochran-Mantel-Haenszel test after FDR correction. Comparisons were between w.t and *pnk-1(tm8231)* mutants per RNAi treatment **(A,B,F,G)** or relative to the corresponding control RNAi sample **(C,E)**.

Next, we examined whether autophagy is implicated in the improvement of proteostasis upon *pnk-1*/ISCs deficiency. First, we examined whether RNAi treatment against *pnk-1* and ISC related genes improves heat shock resistance of autophagy mutants. We found that perturbations in the *pnk-1*/ISCs pathways increased heat shock survival of *atg-4.1* and *atg-9* autophagy mutants (**Figure 3C,S5A)**, as it did in wild-type animals (**Figure 2H)**. Likewise, silencing of *pnk-1* in *atg-4.1* mutants still significantly improved DAF-28::GFP secretion (**Figure 3D).** Finally, only a marginal change in the levels of the autophagy substrate SQST-1::GFP was observed upon *pnk-1* RNAi treatment **(Figure S5B)**.These results indicate that macro-autophagy is not required for proteostasis improvement by *pnk-1*/ISCs deficiency.

Since proteins can undergo lysosomal degradation also through alternative autophagy pathways such as chaperone mediated autophagy and micro-autophagy (Tekirdag & Cuervo 2018), we investigated whether *pnk-1* deficiency-mediated proteostasis improvement implicated the lysosome. To this end, wild-type animals were treated with control RNAi or with RNAi against *pnk-1*/ISC related genes in the presence of 30mM chloroquine (CQ), a lysosomal inhibitor which neutralizes lysosomal acidity and consequently interferes with the degradation activity within the lysosome. The survival of the animals was evaluated 16 hours post heat shock treatment. Despite the decreased survival rate in the CQ-treated groups, deficiency in *pnk-1* and ISC related genes increased animal survival even in the presence of CQ (**Figure 3E)**. Similarly, *pnk-1* mutants survived heat shock better than their wild-type controls, even upon treatment with RNAi targeting components of the lysosomal v-ATPase subunits responsible for lysosomal acidification (**Figure 3F, S5C)**. Thus, *pnk-1* deficiency improves proteostasis independent of the degradative functions of the lysosome.

Since proteostasis improvement by *pnk-1* deficiency is independent of the proteosome and the lysosome, we next explored the possibility that it may rely on improved protein folding rather than protein degradation. Protein refolding is carried out by dedicated chaperone proteins. To test whether chaperone-based protein processing might be required for proteostasis improvement upon reduction of *pnk-1* expression, we screened for the effect of RNAi treatment against a selection of chaperones (14 cytosolic chaperones and 5 ER resident chaperones) on the survival of wild-type and *pnk-1(tm8231)* mutants. Strikingly, RNAi treatment against 9/14 of the cytosolic chaperones and against 4/5 of the ER resident chaperones compromised the relative survival of *pnk-1* mutants compared to the matched RNAi-treated wild-type animals (**Table S3**). We confirmed that RNAi treatment against four of the chaperones identified in the mini-chaperone screen indeed compromised heat-shock survival of *pnk-1* mutants (**Figure 3G**). The finding that many chaperones contribute to the heat shock resistance of *pnk-1* mutants, along with the previous findings that *pnk-1* deficiency improves the proportion of properly folded DAF-28 insulin reporter (**Figure 1C**) and improves the folding of a metastable protein (**Figure 1B**), all implicate chaperone-mediated protein folding as a prime mechanism that improves proteostasis in these mutants.

### HLH-30/TFEB binds to the promoter region of many chaperone genes

HLH-30/TFEB serves as a pivotal transcriptional regulator of autophagy and lysosome biogenesis (Lin et al. 2018; Lapierre et al. 2013). Nevertheless, we noticed that 10/13 of the chaperones that contributed to the heat shock survival of *pnk-1* mutants contained an upstream HLH-30 binding site, as defined by published HLH-30 CHIP-seq data (Gerstein et al. 2010). To further explore the possibility that HLH-30 is a major regulator of chaperone genes, we analyzed the overlap between the complete set of 5103 potential HLH-30 CHIP-seq defined targets (Gerstein et al. 2010; Price et al. 2023), and the complete set of 219 *C. elegans* chaperone and co-chaperone genes (Brehme et al. 2014). We identified 92 overlapping genes among these two gene groups, which is significantly higher than expected by chance (Representation factor: 1.7, associated probability p < 2.002e-08) (**Figure 4A, Table S4)**. Likewise, unbiased gene set enrichment analysis (EasyGSEA, Evitta (Cheng et al. 2021)) of the potential HLH-30 CHIP-seq defined targets revealed enrichment in many protein quality control related gene sets, including chaperones, in addition to lysosome and autophagy-related genes, which are most commonly associated with this transcription factor. Specifically, these included genes related to protein folding, response to heat, and the ER and mitochondria unfolded protein responses (**Figure 4B, Table S5**). Interestingly, the pantothenate and CoA biosynthesis pathway genes, including *pnk-1* itself, were also over-represented in the HLH-30 CHIP gene set, suggesting that the HLH-30 transcription factor modulates CoA biosynthesis.

**Figure 4:**
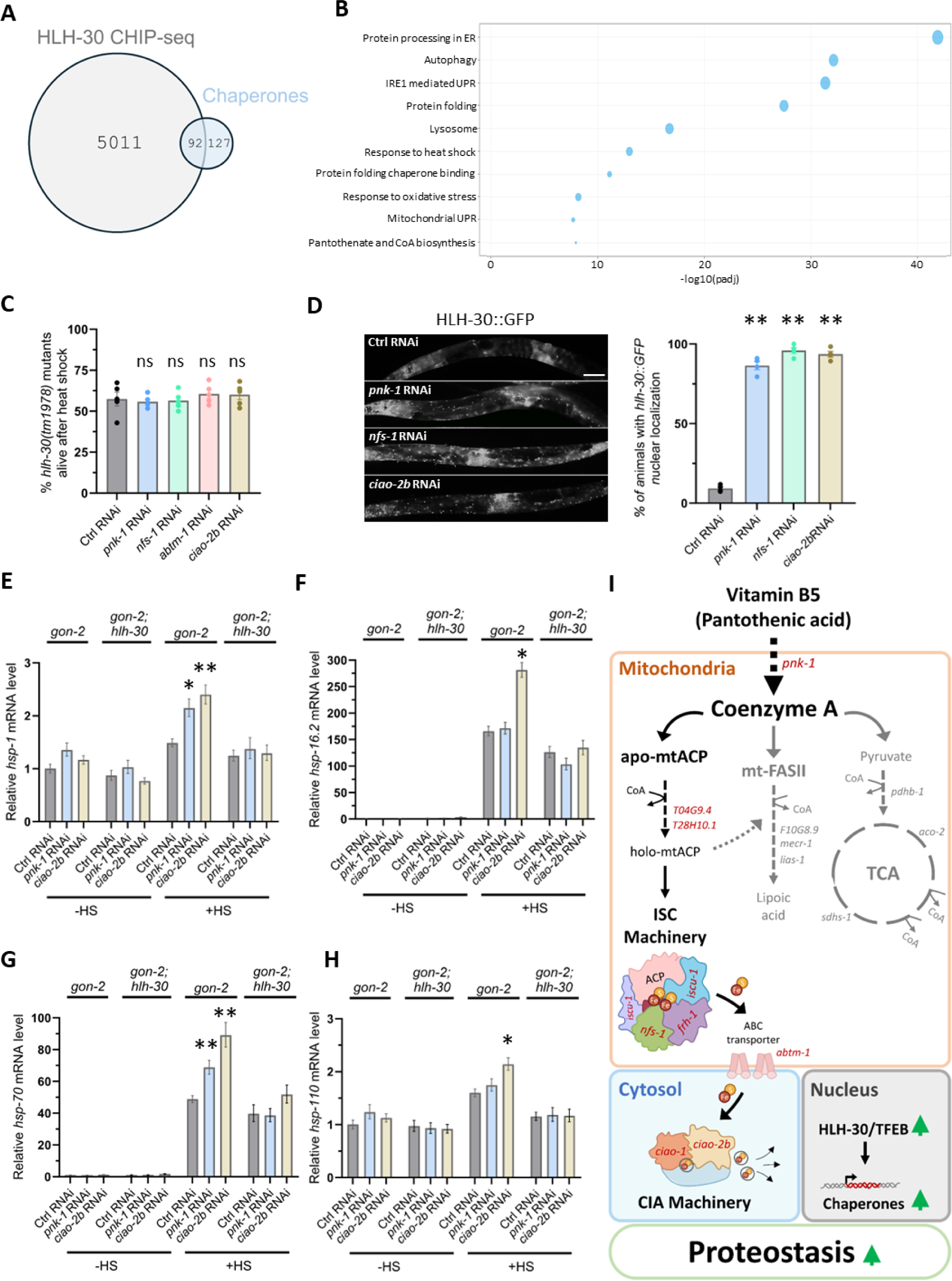
HLH-30 promotes chaperone expression under CoA and ISC mild deficiency. **(A)** Venn diagram depicting an overlap between 5103 potential HLH-30 CHIP-seq defined targets (not including linc genes), and 219 *C. elegans* chaperone and co-chaperone genes (gray and blue, respectively, **Table S4**). Nearly 40% of the chaperone genes overlapped with the group of potential HLH-30 CHIP-seq defined targets. **(B)** Gene set enrichment analysis of the potential HLH-30 CHIP-seq defined targets revealed enrichment in genes related to protein quality control, protein folding, chaperones, response to heat, and the ER and mitochondria unfolded protein responses, lysosome, and autophagy-related genes. Enrichment of pantothenate and CoA biosynthesis genes was also observed. **(C)** *hlh-30(tm1978)* mutation suppressed the benefits of *pnk-1* and ISC related genes inhibition in the heat-shock resistance test. (N=5, n>175). **(D)** *pnk-1* and ISC related genes knockdown increased the nuclear translocation of an HLH-30::GFP transgene in day 3 animals (N=4, n>100). Scale bar: 100μm. **(E-H)** Transcript levels of cytosolic chaperones were examined by qRT-PCR in day 2 adult *gon-2(q388)* and *gon-2(q388); hlh-30(tm1978)* double mutants treated with RNAi against control, *pnk-1* or *ciao-2b* (N=4). In all cases, RNAi treatment against *pnk-1* and/or *ciao-2b* further enhanced the expression of the indicated chaperone genes upon heat shock. **(I)** Summary model: CoA and/or ISC deficiencies improve proteostasis via activation of the conserved HLH-30/TFEB transcription factor. HLH-30/TFEB promotes the expression of a wide range of protein quality control genes, including chaperones, which are critical for the proteostasis improvement. Data are shown as mean ± standard error. N represents the number of biological repeats, n represents the number of animals analyzed per treatment or genetic background. ns, not significant; *P<0.05; **P<0.01. Statistical tests: Cochran-Mantel-Haenszel test followed by FDR correction **(C,D)**, Two Way Anova followed by Tukey’s post-hoc analysis **(E,F,G,H)**. Comparisons were relative to the corresponding control RNAi sample.

### HLH-30/TFEB is required for proteostasis improvement under CoA/ISC limiting conditions

To examine whether a deficiency in the *pnk-1* pathway leads to improved proteostasis via HLH-30/TFEB, we subjected *hlh-30* mutants to *pnk-1* RNAi and assessed their survival under heat shock. Unlike wild-type animals, whose heat shock resistance was improved by RNAi treatments against *pnk-1* or ISC related-genes (**Figure 2H)**, no improvement in survival by these RNAi treatments was observed in *hlh-30* mutants (**Figure 4C).** Additionally, in the DAF-28::GFP secretory assay, the absence of *hlh-30* completely repressed the positive impact of *pnk-1* RNAi **(Figure S6).** Furthermore, increased nuclear translocation of an HLH-30::GFP transgene was observed in day 3 animals exposed to RNAi against *pnk-1* or ISC related genes (**Figure 4D).** These findings demonstrate that the TFEB transcription factor HLH-30 is activated by deficiencies in CoA/ISCs, and its activity is required for improving proteostasis.

### HLH-30/TFEB potentiates chaperone expression in heat-stressed CoA/ISC deficient animals

Since *hlh-30* is important for proteostasis improvement in *pnk-1* mutants and given that this resilience to proteostasis stress in CoA/ISC limiting conditions was chaperone dependent but lysosome independent, we next examined whether transcripts of cytosolic chaperones with putative HLH-30 binding sites were upregulated in response to *pnk-1*/ISC pathway deficiency, and whether this may be mediated by the HLH-30/TFEB transcription factor.

To this end, chaperone transcript levels were examined by qRT-PCR in *gon-2(q388)* and *gon-2(q388); hlh-30(tm1978)* double mutants treated with RNAi against control, *pnk-1* or *ciao-2b* (**Figure 4E-H, Table S6)**. The *gon-2(q388)* temperature sensitive mutation impairs gonad development (Sun & Lambie 1997), ensuring that the detected transcripts represent the stress response of the mothers rather than that of the embryos/germline. Importantly, *gon-2(q388)* mutation had no impact on the heat-shock benefits conferred by the *pnk-1* pathway deficiency **(Figure S7)**. Strikingly, *pnk-1* RNAi enhanced the induction in the transcript levels of 2/4 cytosolic chaperones examined upon heat shock **(Figure 4E, G)**. The heat-shock induction of the transcripts of all four cytosolic chaperones we examined was more pronounced upon treatment with *ciao-2b* RNAi (**Figure 4E-H)**. In the case of *hsp-1* and *hsp-110* genes, the heat shock-induced increase in their transcript levels was only observed upon knockdown of *pnk-1/ciao-2b* (**Figure 4E, H)**. In the case of *hsp-16.2* and *hsp-70* genes, knockdown of *pnk-1/ciao-2b* further increased the levels of their transcripts, beyond the basal induction observed upon heat shock (**Figure 4F, G)**. In all cases, *pnk-1/ciao-2b* RNAi mediated increase in chaperone transcript levels was only observed upon heat stress and was *hlh-30*-dependent **(Figure 4E-H, Table S6)**. This suggests that HLH-30 boosts the stress-induced transcription of cytosolic chaperones under CoA/ISC limiting conditions.

## Discussion

Loss of function mutations in PANK limit CoA biosynthesis and are usually associated with neurodegenerative disorders such as Pantothenate kinase associated neurodegeneration (PKAN), dystonia, Parkinson’s disease, and Alzheimer’s disease (Kruer et al. 2011; Hayflick et al. 2003; Brunetti et al. 2012; Burté et al. 2015). However, less is known about the consequences of mild deficiencies in *PANK* homologs.

In this study we employed the model organism *C. elegans* to investigate the consequences of mild deficiency in *pnk-1*. Unlike the characteristic pathology of the *pnk-1* knockout mutant, hypomorphic *pnk-1* mutants had a normal lifespan and were resistant to a wide range of proteostasis challenges, in the ER and in the cytosol. Importantly, inhibition of PANK protected from proteostasis-related stress also in a human cell line system, demonstrating the conservation and cell-autonomous nature of this proteostasis promoting pathway. Proteostasis improvement by *pnk-1* knock down was suppressed by external supplementation with CoA, indicating that limitation of CoA biosynthesis triggers the observed proteostasis improvement. Together, this suggests that CoA limitation can be beneficial under conditions that challenge proteostasis.

Among the many CoA-dependent events in the mitochondria (Leonardi et al. 2005), knockdown of the ISC machinery also had a robust proteostasis improving effect. Epistasis experiments indicated that *pnk-1* deficiency and ISC related genes knockdown both improve proteostasis by a common mechanism. This suggests that CoA limitation promotes proteostasis by limiting ISC formation, which in turn enhances proteostasis. Interestingly, feeding *C. elegans* bacteria with aberrant Fe-S cluster biogenesis also enhances protein stress tolerance of the *C. elegans* host (Bhat et al. 2024), suggesting that ISC deficiency can be controlled through the diet. Since knockdown of the ISC transporter responsible for exporting ISCs from the mitochondria to the cytosol and knock down of proteins that load the ISCs onto target cytosolic Fe-S proteins also improved proteostasis, this uncouples between mitochondrial and non-mitochondrial Fe-S related functions. Hence, they implicate cytosolic/nuclear Fe-S proteins rather than mitochondrial Fe-S proteins in the regulation of proteostasis.

At the mechanistic level, proteostasis improvement by CoA/ISC deficiencies are dependent on chaperones, yet independent of protein quality control mechanisms related to protein degradation such as the proteasome and the lysosome. This suggests that the chaperones that promote proteostasis upon CoA/ISC depletion do so by supporting the refolding of misfolded proteins, by solubilizing aggregated proteins or preventing their aggregation, rather than by clearing these aberrant proteins via the cellular degradation machineries (Kim et al. 2013). Interestingly, in spite the fact that hundreds of chaperones are encoded in the genome, the improved resistance to proteostasis challenges was lost upon silencing of individual chaperones. This may reflect the lack of redundancy among the various chaperones, which may have different substrate preferences, and thus support different client proteins, in the context of proteostasis challenges.

Proteostasis improvement by CoA/ISC deficiencies is also dependent on the transcription factor HLH-30/TFEB, which translocates to the nucleus upon CoA/ISC depletion. Interestingly, the CoA biosynthesis pathway itself is over-represented among the HLH-30 target genes, and TFEB-dependent increase in CoA biosynthesis has been reported in mice in response to sulfur amino acid metabolism (Matye et al. 2022), constituting a conserved negative feedback loop between sulfur and CoA availability and TFEB.

HLH-30/TFEB is best known for its crucial role in lysosome biogenesis and autophagy (Franco-Juárez et al. 2022; Lin et al. 2018; Wong et al. 2023; Settembre et al. 2011), and as an essential transcription factor required and sufficient for most longevity pathways in *C. elegans* (Lapierre et al. 2013). Nevertheless, neither lysosome biogenesis nor autophagy are required for the improved proteostasis upon CoA/ISC depletion. Instead, the improved proteostasis relies on a chaperone-mediated proteostasis shield, mediated by individual chaperones whose transcription is potentiated under proteostasis-challenging conditions in an HLH-30-dependent manner. Indeed, analysis of HLH-30 target genes identified in ChIP experiments reveals that many proteins quality control related gene sets, including chaperones and unfolded protein response genes alongside the classical lysosome and autophagy-related genes, are enriched within the potential target genes of the HLH-30 transcription factor. This raises the notion that the pro-longevity (Lapierre et al. 2013) and pro-proteostasis (Wong et al. 2023; Lin et al. 2018) properties of HLH-30 may be attributed to the rewiring of the entire protein quality control network. Importantly, human TFEB has also been identified as a component of the integrated stress response (Martina et al. 2016), demonstrating its importance beyond the autophagy and the lysosomal pathways.

Altogether, HLH-30/TFEB controls a wide-range of protein quality control mechanisms, implying it is an evolutionary conserved key longevity and proteostasis-promoting transcription factor, which induces the expression of chaperone and stress response genes alongside autophagy/lysosome genes, all of which may contribute to improved proteostasis and health span. These results highlight TFEB and its upstream regulators as potential therapeutic targets in proteostasis-related diseases.

## Methods

### *C. elegans* strains used in this study

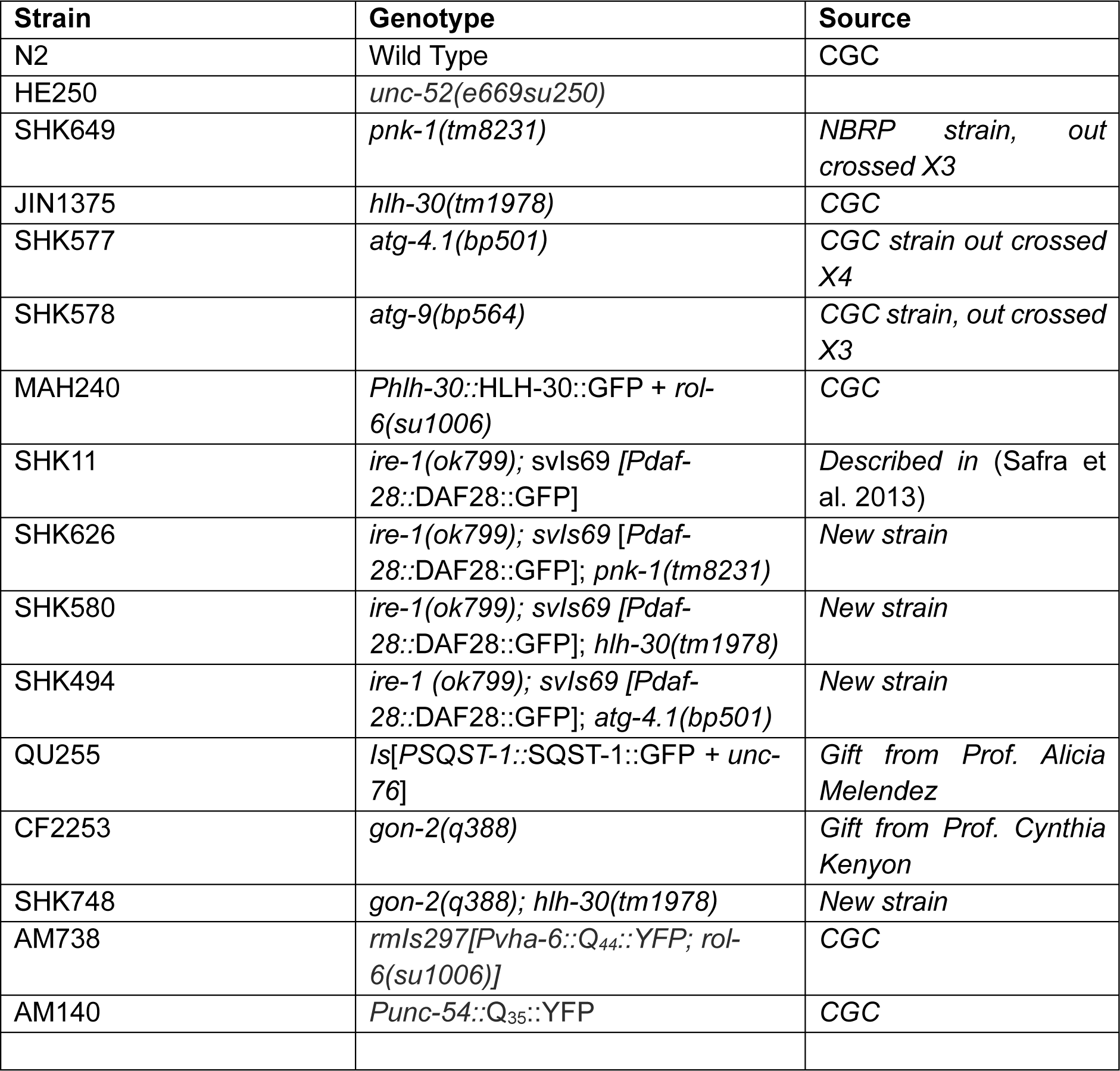

### Worm maintenance

Strains were maintained at 20°C using standard *C. elegans* methods. Nematodes Growth Media (NGM) agar plates were seeded with *E. coli* strain OP50 or with HT115 bacteria for RNAi experiments. For CoA supplementation experiment, regular NGM plates served as a control and treatment plates were supplemented with 400μM Coenzyme A (Sigma-Aldrich, C4780).

### RNA interference

HT115 bacteria expressing dsRNA were cultured overnight in LB containing 10μg/mL tetracycline and 100μg/mL ampicillin. Before seeding, IPTG was added to the bacteria medium to a final concentration of 20μM. 500μl bacteria were seeded on NGM plates containing 2mM IPTG and 0.05mg/ml carbenicillin. RNAi clone identity was verified by sequencing. Eggs were placed on fresh plates (2 days after bacteria seeding) and synchronized at L4. Animals were fed with RNAi bacteria starting from eggs and throughout the experiment, except for *rpn-6.1*, *vha-12* and *vha-1* RNAi treatments in which animals were grown on control RNAi plates from eggs till L4stage and transferred to plates with these RNAi bacteria starting from L4 stage.

### DAF-28::GFP secretion assay

Eggs of *ire-1(ok799)* animals expressing a *Pdaf-28::daf-28::gfp* transgene were grown at 20°C on NGM or RNAi plates until day 3 of adulthood. 3 day old transgenic animals were anaesthetized on 2% agarose pads containing 2 mM levamisole (Sigma, L9756). DAF-28::GFP expression pattern was examined in at least 30 animals on day 3 of adulthood. Animals in which the transgene was detected within the body cavity of the animals were scored as animals with improved DAF-28::GFP secretion. Animals in which the transgene was trapped within the producing cells in the posterior intestine were scored as animals with a secretory defect.

### Thrashing test

*unc-52(e669su250)* eggs were incubated at 20°C until L4. Age-synchronized L4 *unc-52(e669su250)* animals were shifted to 25°C until day 2 of adulthood. On day 2 of adulthood, animals were placed in 96 wells containing M9 buffer. After 10 seconds of adjustment period, each animal was monitored and scored for trashing over a 20 second period. Values are presented as body bends per minute.

### Tunicamycin resistance assay

Eggs were placed on fresh plates containing DMSO or 5μM Tunicamycin (Sigma-Aldrich, 654380). After three days, the number of animals that developed to L4 stage/adult stage was scored.

### Heat shock survival assay

Age-synchronized animals were grown at 25°C until day 2 of adulthood. On day 2 of adulthood, animals were subjected to 37°C heat shock for 5.5 hours and recovered at 25°C for overnight (∼16 hours). Animals that failed to move in response to a gentle touch with a metal pick were scored as dead.

For CoA external supplement experiment eggs were grown at 25°C until day 2 of adulthood on regular NGM plates which served as a control or on plates supplemented with 400μM Coenzyme A (Sigma-Aldrich, C4780).

### Chloroquine treatment

Eggs were grown at 25°C until day 1 of adulthood. On day 1 of adulthood, animals were transferred to plates containing 30μM Chloroquine Diphosphate Salt (Sigma, C6628), or maintained on regular plates as controls. On day 2 of adulthood, animals were subjected to 37°C heat shock for 5.5 hours and recovered at 25°C for overnight. Animals that failed to move in response to a gentle touch with a metal pick were scored as dead.

### qRT-PCR

To measure the relative levels of *pnk-1* transcript, w.t and *pnk-1(tm8231)* eggs were raised on NGM plates at 20°C until day3 of adulthood. On day 3 of adulthood synchronized animals were collected for RNA extraction.

To measure the relative levels of select chaperone transcripts, w.t or *hlh-30 (tm1978)* animals were raised on control, *pnk-1* or *ciao-2b* RNAi at 25°C from eggs until day 2 of adulthood. On day 2, animals were exposed to 37°C heat shock for 1 hour and recovered at 25°C for 1 hour. Then, animals were collected for RNA extraction. Total RNA was extracted with TRIzol reagent (Ambion, 15596026) and treated with RNase-free DNase I (Thermo scientific, EN0521). Reverse transcriptions were carried out using the qScript cDNA Synthesis Kit (Quantabio 66225515). Real-time PCR was done using Blue Mix Hi-Rox SYBR GREEN (PCR Biosystems, Pb20-16-05) in a StepOnePlus™ Real-Time PCR System. Transcript levels were analyzed by the ΔΔCT method. Transcript levels of *ama-1* were used for normalization. Each sample was run in triplicates, and four independent biological samples were analyzed. P-values were calculated using One Sample T-test for *pnk-1* mRNA. For chaperones transcripts P-values were calculated using Two Way Anova followed by Tukey Post Hoc.

Primers used for qPCR:

1. *pnk-1:* FW: 5’-GGACAGATGGCATATTTGTATGG -3’ BW: 5’-CAATCTCTCCCTTGCTCCAATA -3’
2. *hsp-1:* FW: 5’-GTTGCCATGAACCCACATAAC -3’ BW: 5’-CGGCAGAGATGACCTTGAAT -3’
3. *hsp-16.2:* FW: 5’-CTGAGTCTTCTGAGATTGTT -3’ BW: 5’-CGTACGACCATCCAAATTA -3’
4. *hsp-70:* FW: 5’-CAATGGGAAGGACCTCAAC -3’ BW: 5’-GGGACAACATCAACGAGTAA -3’
5. *hsp-110:* FW: 5’-CAGGCTTCATTGGTTGCT -3’ BW: 5’-GCGGCATCAATTCCGTAT -3’
6. *ama-1:* FW: 5’-CGGAGGAGATTAAACGCATGT -3’ BW: 5’-GCATCTTCCACGACGATCTATG -3’

### Lifespan

Eggs were placed on plates seeded with the RNAi bacteria of interest and were continuously transferred to freshly seeded RNAi plates. Lifespan was scored every 1–2 days. Related lifespans were performed concurrently to minimize variability. In all experiments, lifespan was scored as of the L4 stage which was set as t = 0. Animals that ruptured or crawled off the plates were included in the lifespan analysis as censored worms. SPSS program was used to determine the means and the P values. P values were calculated using the Log Rank (Mantel-Cox) and Breslow (Generalized Wilcoxon) method.

### Western blot

100 day-3 animals were boiled in protein sample buffer containing 2% SDS. Proteins were separated using standard PAGE separation, transferred to a nitrocellulose membrane, and detected by western blotting using anti-tubulin (Sigma, T5168, 1∶1000) and anti-GFP (Roche, 11814460001, 1:1,000) antibodies.

### Fluorescence microscopy and quantification

To follow expression of fluorescent transgenic markers, animals were anaesthetized on 2% agarose pads containing 2 mM levamisole. Images were taken with a CCD digital camera using a Nikon 90i fluorescence microscope. For each trial, exposure time was calibrated to minimize the number of saturated pixels and was kept constant through the experiment unless indicated otherwise. The NIS element software was used to quantify mean or sum fluorescence intensity of manually selected regions of interest encompassing individual animals.

### Cell culture

Human U2OS cells (ATCC) were maintained in low glucose DMEM (Biological Industries, Beit-Haemek, Israel) supplemented with 10% fetal bovine serum (FBS; HyClone Laboratories, Logan, UT). Cells were seeded in 6 well plates to a density of 60-70%. After 24 hours, cells were treated with PANK inhibitor (250nM or 500nM) or with DMSO as a control. 16 hours following the treatment, cells were exposed to 2 hours heat shock at 42-43 degrees. Immediately thereafter, double staining with Annexin V-FITC and PI was performed using an Annexin V-FITC Apoptosis Kit (Bio Vision K101-100-3). Flow cytometry analysis was performed on a BD LSR Fortessa Cell Analyzer. Cell viability was determined by the population negative for annexin V and PI.

### Statistical Analysis

All experiments results are presented as the mean of at least three independent experiments. All error bars show the SEM, unless otherwise stated. Statistical significance was determined by using statistical tests as indicated in the figure legends and in Table S7. Statistical P-values of <0.05 were considered significant. Data were analyzed using SPSS. Graphs were created by GraphPad Prism software.

## Supporting information

Tasble S1

Tasble S2

Tasble S3

Tasble S4

Tasble S5

Tasble S6

Tasble S7

## Data availability

All relevant data can be found within the article and its supplementary information.

## Author Contributions

Conceptualization: R.S., S.H.-K.; Methodology: R.S., M.L.-F., S.H.-K.; Investigation: R.S., M.L.-F., H.C., M.K.H.; Writing - original draft: R.S., M.L.-F., Y.S.-T., S.H.-K.; Supervision: Y.S.-T, S.H.-K.; Funding acquisition: S.H.-K.

## Acknowledgments

We thank members of the Korenblit lab for helpful discussions. We thank Tal Partoosh for preliminary results. We thank Dr. Tamar Juven-Gershon, Hadar Krap Shachar and Dr. Hagit Hauschner for technical assistance. Some nematode strains were provided by the Caenorhabditis Genetics Center, which is funded by the NIH National Center for Research Resources and by Dr. Shohei Mitani, National Bioresource Project for the nematode, Tokyo Women’s Medical University School of Medicine, Japan.

## Figure Legends

**Figure S1:**
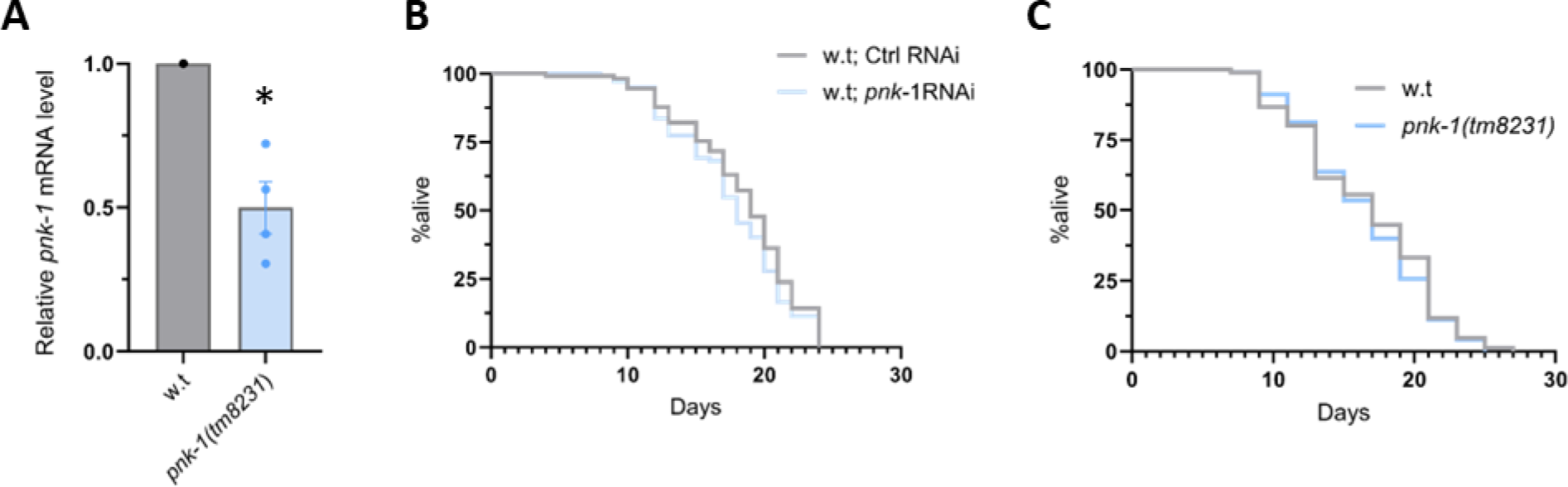
Mild CoA deficiency does not extend lifespan of wild-type animals. **(A)** *tm8231* mutation is a 73 bp deletion ∼3200 bp upstream of the *pnk-1* gene transcription start site. It reduces *pnk-1* transcript levels by nearly 50% as determined by qRT-PCR (N=4). Asterisks mark one sample t-test values of p<0.05. Data are shown as mean ± standard error **(B-C)** Representative lifespan of *pnk-1* hypomorphic mutants. *pnk-1* RNAi treatment **(B)** and the *pnk-1(tm8231)* mutation **(C)** did not extend the lifespan of wild-type animals (N=3). Note that the data of the RNAi lifespan experiment is part of the experiment detailed in table S1.

**Figure S2:**
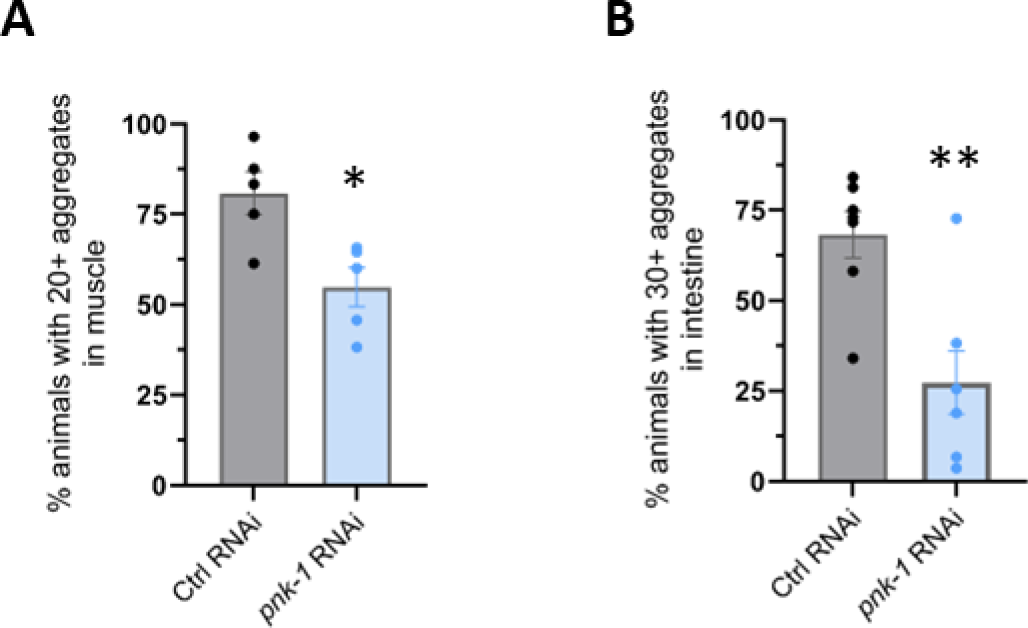
Mild CoA deficiency reduces Poly-Q aggregates. *pnk-1* RNAi reduced the amount of visible aggregates in animals expressing muscular PolyQ35::YFP (N=5, n>125) **(A)** or intestinal PolyQ44::YFP (N=7, n>250) **(B)** in day 5 animals. *P<0.05; **P<0.01. Statistical test: Cochran-Mantel-Haenszel test. Data are expressed as mean ± standard error.

**Figure S3:**
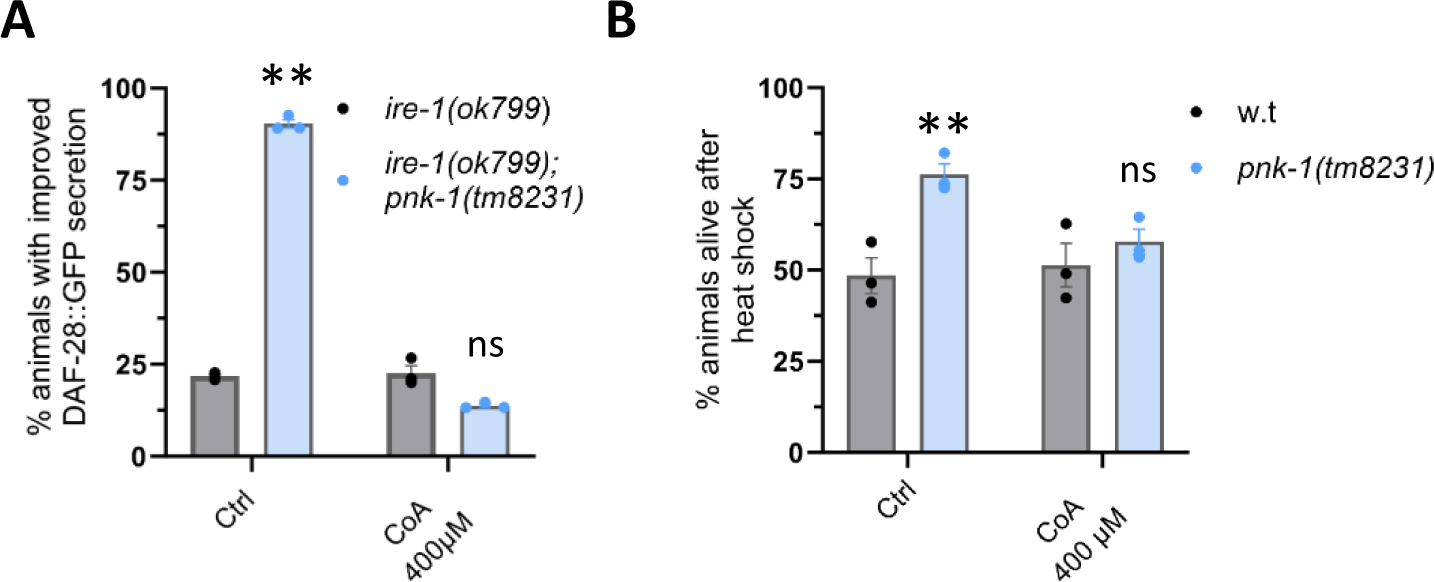
CoA supplementation restores heat shock sensitivity of *pnk-1* mutants. CoA supplementation suppressed the improved secretion of DAF-28::GFP in day 3 *ire-1(ok799); pnk-1(tm8231)* mutants (N=3, n>70) **(A)**, as well as the improved heat-shock resistance of day 2 *pnk-1(tm8231)* mutants (N=3, n>150) **(B)**. ns, not significant; **P<0.01. Statistical test: Cochran-Mantel-Haenszel test followed by FDR correction. Comparisons were between w.t and *pnk-1(tm8231)* mutants per treatment.

**Figure S4:**
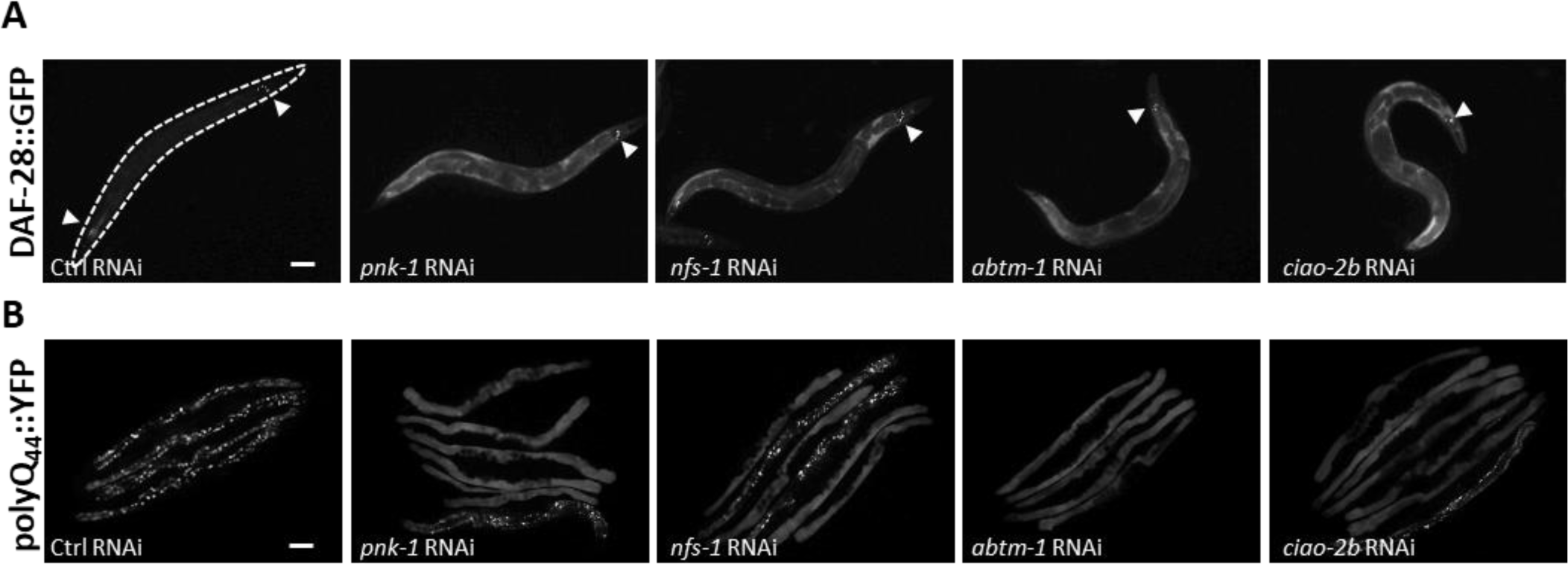
*pnk-1* and ISC related genes RNAi improve protein secretion and reduce aggregates. **(A)** Representative images of DAF-28::GFP in the *ire-1(ok799)* mutants treated with *pnk-1* and ISC related genes RNAi. Arrow heads mark head neurons and intestinal cells that accumulate DAF-28::GFP protein. Knockdown of *pnk-1*, ISC (*nfs-1*) and CIA (*abtm-1, ciao-2b*) related genes improved the secretion of DAF-28::GFP. Scale bar: 100μm. **(B)** Representative images of intestinal PolyQ44:YFP aggregates in day 5 adult animals treated with *pnk-1* and ISC related genes RNAi. Scale bar: 100μm.

**Figure S5:**
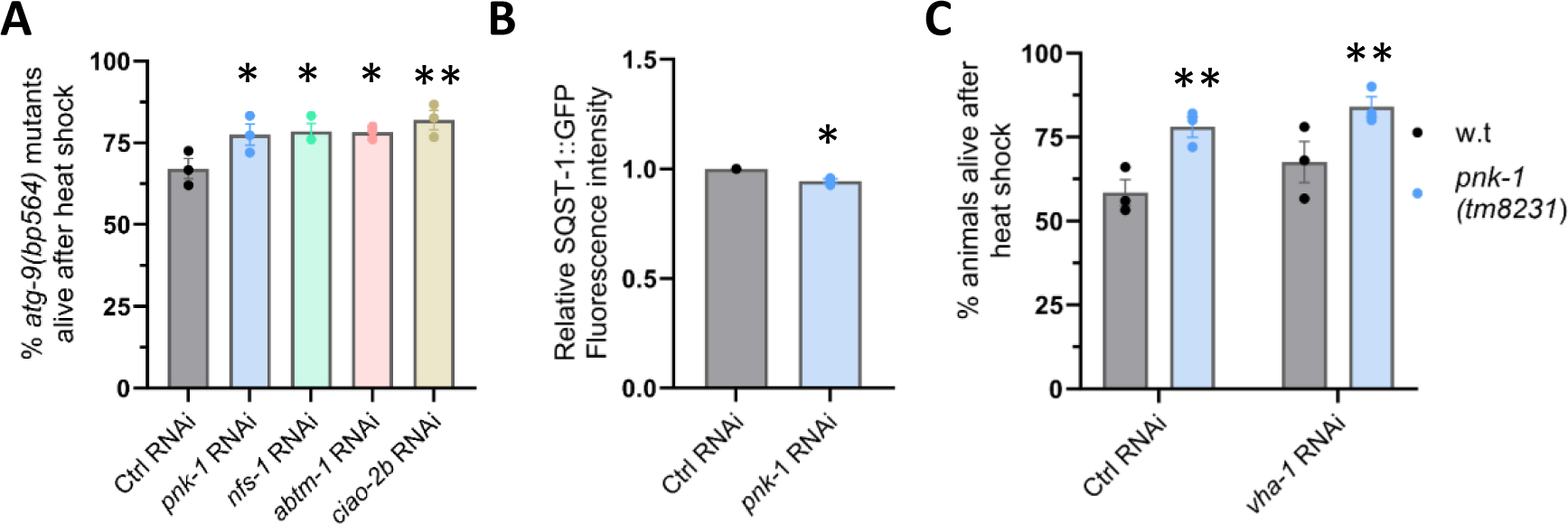
*pnk-1* and ISC related genes inhibition promote proteostasis independent of the autophagy-lysosome system. **(A)** RNAi inhibition of *pnk-1* and ISC related genes improved heat-shock resistance in *atg-9(bp564)* mutants. (N=3, n>90). **(B)** *pnk-1* RNAi resulted in a very minor decrease in the level of the autophagy substrate reporter SQST-1::GFP. (N=3, n>90). **(C)** Inhibition of lysosome function by *vha-1* RNAi treatment in adult animals did not affect heat-shock resistance of *pnk-1(tm8231)* animals. (N=3, n>90). Data are shown as mean ± standard error. N represents the number of biological repeats, n represents the number of animals analyzed per treatment or genetic background. *P<0.05; **P<0.01. Statistical tests: one sample t-test **(B)**; Cochran-Mantel-Haenszel test **(A,C)**. Comparisons were between w.t and *pnk-1(tm8231)* mutants per treatment **(C)** or relative to the corresponding control RNAi sample **(A,B)**.

**Figure S6:**
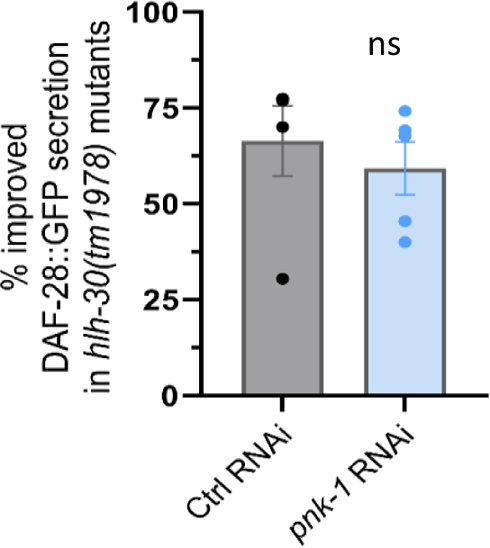
*hlh-30* transcription factor is required for properly ER protein secretion upon *pnk*-1 mild deficiency in ire-1 mutants. *hlh-30(tm1978)* mutation prevented the benefits of *pnk-1* RNAi in the DAF-28::GFP secretion test. (N=5, n>110). ns marks Cochran-Mantel-Haenszel test values of p>0.05. ata are shown as mean ± standard error. N represents the number of biological repeats, n represents the number of animals analyzed per RNAi treatment.

**Figure S7:**
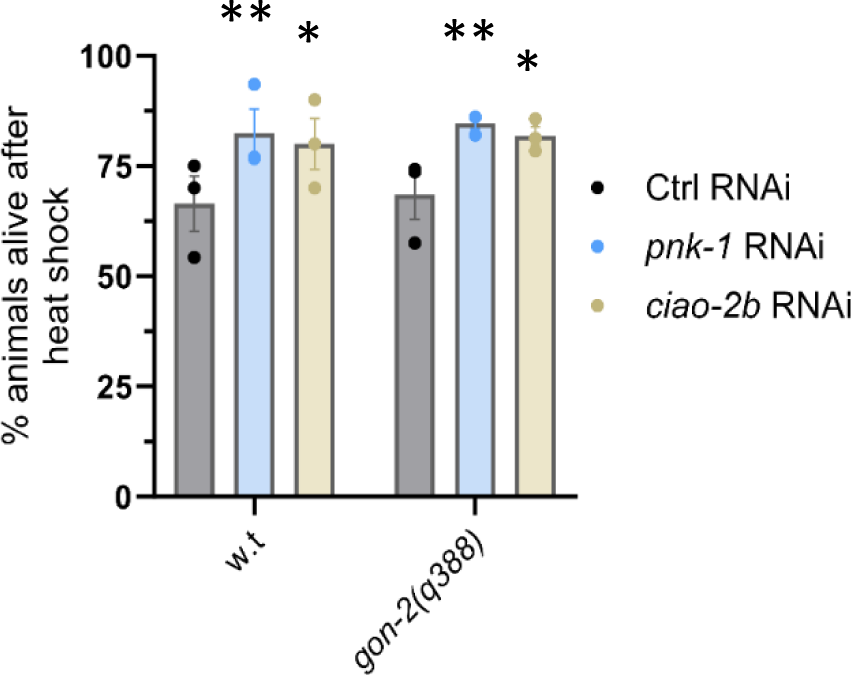
CoA and ISC mild deficiency improves proteostasis in *gon-2* mutants. *pnk-1* and *ciao-2b* RNAi treatments improve the heat shock resistance of *gon-2(q388)* gonadless animals (N=3, n>90), as it does in w.t animals. Data are shown as mean ± standard error. N represents the number of biological repeats, n represents the number of animals analyzed per treatment or genetic background. *P<0.05; **P<0.01. Statistical tests: one sample t-test. Comparisons were relative to the corresponding control RNAi sample.

## Supplementary Tables

**Table S1: The effect of pnk-1/ISCs deficiency on lifespan in C. elegans.**

**Table S2: Inhibition of PANK activity promotes heat shock resistance in human osteosarcoma U2OS cells.**

**Table S3: chaperone RNAi screen.**

**Table S4: Data sets of potential HLH-30 CHIP-seq defined targets and 219 C. elegans chaperone and co-chaperone gene.**

**Table S5: Gene set enrichment analysis of the potential HLH-30 CHIP-seq defined targets.**

**Table S6: Cytosolic chaperones qRT-PCR data.**

**Table S7: Statistics.**

